# First case of three-way multiple herbicide resistance in *Amaranthus palmeri* outside the Americas: Resistance mechanisms and alternative management strategies

**DOI:** 10.64898/2026.09.07.749782

**Authors:** Maor Matzrafi, Jackline Abu-Nassar, Amit Paporisch

## Abstract

The rapid spread of herbicide-resistant *Amaranthus palmeri* S. Watson across the Mediterranean region poses a severe threat to crop production. This study characterizes the resistance profile of an *A. palmeri* population from Gonen, Israel, which survived multiple post-emergence herbicide applications in maize. Greenhouse dose-response and screening trials were conducted across four modes of action (MOAs). The Gonen population exhibited three-way multiple resistance to acetolactate synthase (ALS), photosystem II (PSII) and 4- hydroxyphenylpyruvate dioxygenase (HPPD) inhibitors, marking the first confirmed case outside the Americas. High-level resistance was demonstrated for the ALS inhibitor foramsulfuron and the PSII inhibitor metribuzin, maintaining 40-70% and 80-100% survival, respectively, at double the labelled field rate. Sequencing of the ALS gene revealed a Trp574 to Leu substitution in resistant plants. The population showed differential PSII sensitivity, as atrazine provided complete control despite high metribuzin survival. Sequencing revealed no known *psbA* mutations, and malathion pre-treatment caused only a slight reduction in metribuzin survival. Partial resistance was observed for the HPPD inhibitor tembotrione (30- 50% survival), where malathion pre-treatment reduced survival by 20-30%. In contrast, pre-mergence applications of *S*-metolachlor (very long chain fatty acid inhibitor), isoxaflutole (HPPD inhibitor), and a fomesafen + terbutryn mixture [protoporphyrinogen oxidase (PPO) + PSII inhibitors] significantly reduced emergence. These findings confirm multiple-resistant *A. palmeri* in the region and highlight the urgent need for integrated weed management incorporating effective pre-emergence herbicides rather than relying solely on post-emergence chemical control in summer cereal rotations.

## 1 INTRODUCTION

*Amaranthus palmeri* S. Watson (Palmer amaranth) is an aggressive, dioecious summer annual broadleaf weed that poses a major threat to agricultural productivity worldwide (Roberts & Florentine, 2022). Characterized by exceptionally rapid vegetative growth, high fecundity, and high genetic diversity, *A. palmeri* rapidly establishes dominance in disturbed agroecosystems, outcompeting major field crops for light, nutrients, and moisture (Ward et al., 2013). Native to the southwestern United States and northern Mexico, *A. palmeri* has expanded its geographic range across North America and, in recent decades, invaded the Mediterranean region and multiple European countries (Torra et al., 2020; Matzrafi et al., 2023). The species’ evolutionary plasticity has facilitated its rapid adaptation to diverse climatic conditions and intensive agricultural practices, making it an invasive weed of international concern.

The management of *A. palmeri* has become increasingly complex due to its propensity to evolve resistance to multiple herbicide modes of action (MOAs). Biotypes resistant to acetolactate synthase (ALS - HRAC 2) inhibitors, synthetic auxins, photosystem II (PSII - HRAC 5) inhibitors, 4-hydroxyphenylpyruvate dioxygenase (HPPD - HRAC 27) inhibitors, 5-enolpyruvylshikimate-3-phosphate synthase (EPSPS - HRAC 5) inhibitors, and protoporphyrinogen oxidase (PPO - HRAC 14) inhibitors have been documented globally, with several populations exhibiting complex multiple-herbicide resistance (Gaines et al., 2020; Heap, 2026). In the Mediterranean basin, populations with resistance to various MOAs have recently been reported in Turkey, Greece, Spain, and Italy, underscoring the growing systemic threat to European crop production (Torra et al., 2020; Kanatas et al., 2021; Mennan et al., 2021; Milani et al., 2021). In Israel, *A. palmeri* has spread extensively throughout major cropping regions, where ALS-inhibitor resistance is already widespread (Matzrafi et al., 2024).

Chemical weed management in Israeli summer cereal systems, particularly maize (*Zea mays* L.), depends on a narrow spectrum of active ingredients. Pre-emergence weed control relies largely on the very-long-chain fatty acid (VLCFA - HRAC 15) inhibitor *S*-metolachlor and Isoxaflutole (HPPD), whereas post-emergence programs rely on a restricted set of mode-of-action groups, including ALS inhibitors (e.g., foramsulfuron), PSII inhibitors (e.g., atrazine), HPPD inhibitors (e.g., tembotrione), and synthetic auxins (e.g., fluroxypyr). However, recent field observations in maize-growing regions have indicated a sharp decline in herbicide efficacy, especially for post-emergence products, resulting in weed control failures and substantial yield losses. Reduced herbicide performance in the field can stem from multiple factors, including environmental degradation of soil-applied herbicides, improper application timing, or the selection and proliferation of herbicide-resistant biotypes (Chauhan, 2020).

Understanding whether control failures are driven by target-site mutations or non-target-site resistance (NTSR) mechanisms, such as enhanced herbicide detoxification mediated by cytochrome P450 monooxygenases, is critical for designing sustainable weed management strategies (Powles & Yu, 2010; Gaines et al., 2020).

To address these concerns, this study investigated a suspicious *A. palmeri* population collected from a maize field near Kibbutz Gonen, where commercial herbicide programs consistently failed. The primary objectives of this study were to: (1) evaluate the baseline sensitivity and resistance profile of the Gonen population across four major herbicide MOAs (ALS, PSII, HPPD, and auxin mimics) across a range of doses in comparison to a known susceptible population (GE’A); (2) elucidate potential resistance mechanisms; and (3) evaluate pre-emergence herbicides to identify alternative chemical control options for integrated weed management.

## 2 MATERIALS AND METHODS

### 2.1 Seed collection method

*Amaranthus palmeri* seeds of the suspected herbicide-resistant population were collected at a maize field near kibbutz Gonen (33°07’30.3”N, 35°38’06.7”E), using a sampling technique previously described by Culpepper et al. (2008) with minor adjustments. Seed collection includes randomly harvesting 4-6 seed heads from each plant for approximately 40 female plants spaced at least 10 m apart, collated haphazardly over the field. Seed heads were threshed and stored at 4°C until use. Seeds from the sensitive GE’A population, previously described by Abu-Nassar et al. (2025), were used to determine the baseline level of resistance.

### 2.2 Herbicide application

#### 2.2.1 Pre-emergence herbicide application

To evaluate efficacy of herbicides of pre-emergence herbicides, a controlled-condition pot experiment was conducted. For this study, we used a 50 L composite soil sample, from the top 10 cm soil layer of the Gonen field. Soil was sampled from at least 10 different points at the edges of the field, an area to which residual herbicides were not applied for at least one year prior to sampling. The soil was packed into 0.3 L pots and sown with *A. palmeri* seeds from Gonen population. Following sowing, herbicides were applied as specified in Table 1. The experiment utilized 10 replicates for each herbicide treatment and control. Pots were sprayed using a chain-driven sprayer delivering 300 L ha^-1^ with a flat-fan 8001E nozzle at 2 bars (TeeJet®, Spraying Systems Co., Wheaton, IL, USA). Experiments were conducted at Newe-Ya’ar Research Center in a greenhouse with at an average temperature of 33/22±3°C day/night and a photoperiod of 13-14 hours. Pots were placed on experimental benches and randomized every week. After application, the pots were overhead-irrigated to incorporate the herbicides into the soil solution and then placed in a greenhouse for approximately two months, with daily bottom irrigation, to avoid leaching of herbicides. At 29 days after treatment (DAT), the number of emerged *A. palmeri* plants was recorded, and the shoots were harvested to measure fresh weight. Subsequently, the top layer of soil in each pot was lightly tilled, and a second cohort of *A. palmeri* seeds was sown. This second cohort was grown for an additional 22 days at the same conditions (totaling 51 DAT), after which plant count and fresh weight were again recorded.

**Table 1.**
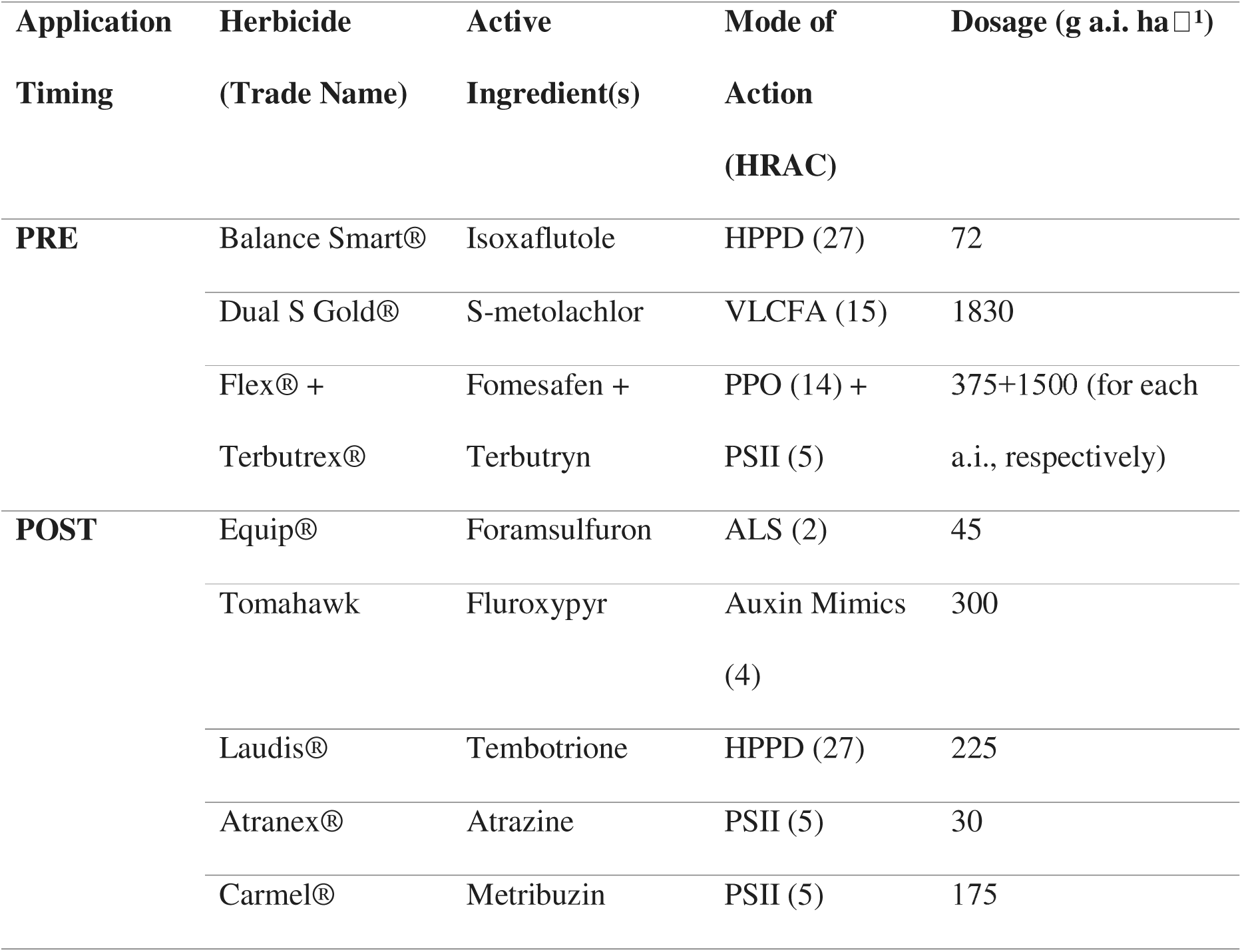
Herbicides evaluated for the control of *A. palmeri* in pre-emergence (PRE) and post-emergence (POST) greenhouse trials.

#### 2.2.2 Post-emergence herbicide application

Seeds were sown in 0.3L pots filled with a commercial growth mixture (Tuf Marom-Golan®). Upon emergence, seedlings were thinned to obtain one seedling per pot. When plants reached the 5-7 cm tall growth stage (4-6 leaf stage), they were sprayed using the recommended labelled field rate (Table 1). Plants were sprayed and maintained under the same conditions as described above. Shoot fresh weight and percentage survival was recorded 21 DAT. Survival was defined as the ability to produce new leaves for a treated individual. The experiment was conducted twice with 10 replicates for each experimental run.

### 2.3 Mechanism of resistance

#### 2.3.1 Malathion treatment

Seeds from both populations (Gonen and GE’A) were sown in pots (as detailed above), and seedlings were thinned following emergence to a single plant per pot. When plants reached the 5-7 cm tall growth stage (4-6 leaf stage), they were sprayed using the recommended labelled field rate of tembotrione and metribuzin, either solely or preceded by a treatment of malathion at a rate of 1000 g ha^-1^ applied two hours prior to herbicide application. Control treatments included malathion-only application and untreated control plants. Shoot fresh weight and percentage survival was recorded 21 DAT. Survival was defined as the ability to produce new leaves for a treated individual. The experiment was conducted twice with 10 replicates for each experimental run.

#### 2.3.2 Sequencing of the *ALS* and PSII genes

In order to detect structural substitutions, the *ALS* and *psbA* genes were sequenced and analysed. A leaf tissue was excised from each plant that have survived the treatment with double the recommended labelled field rate of with either foramsulfuron or metribuzin from Exp. 2 (seven and eight replicates, respectively). DNA extraction was performed using the same procedures as described in Matzrafi et al. (2017). Amplification and sequencing of the *psbA* and *ALS* genes followed PCR protocols and primer sets from Sibony et al. (2003) and Reinhardt et al. (2022), respectively. Sequences were aligned in BioEdit (Hall, 1999) and compared with NCBI reference sequences for *A. retroflexus psbA* (DQ887374.1) and *A. palmeri ALS* (MW361339.1).

### 2.4 Statistical analyses

Figures 1 was generated using RStudio version 4.2.3 (R Core Team, 2023). Differences in plant density and shoot fresh weight between pre-emergence herbicide treatments and the untreated control (UTC) were evaluated separately at 29 and 51 DAT using a One-Way Analysis of Variance (ANOVA). Where global significance was observed (*P* < 0.05), Dunnett’s post hoc test was applied to compare each herbicide regimen against the UTC control baseline. For post-emergence trials, pooling of experimental runs was evaluated using a two-way ANOVA followed by Tukey’s *post hoc* test. Due to significant run-by-treatment interactions between the two repeated trials (*P* < 0.05), Experiments 1 and 2 are displayed independently.

**Figure 1.**
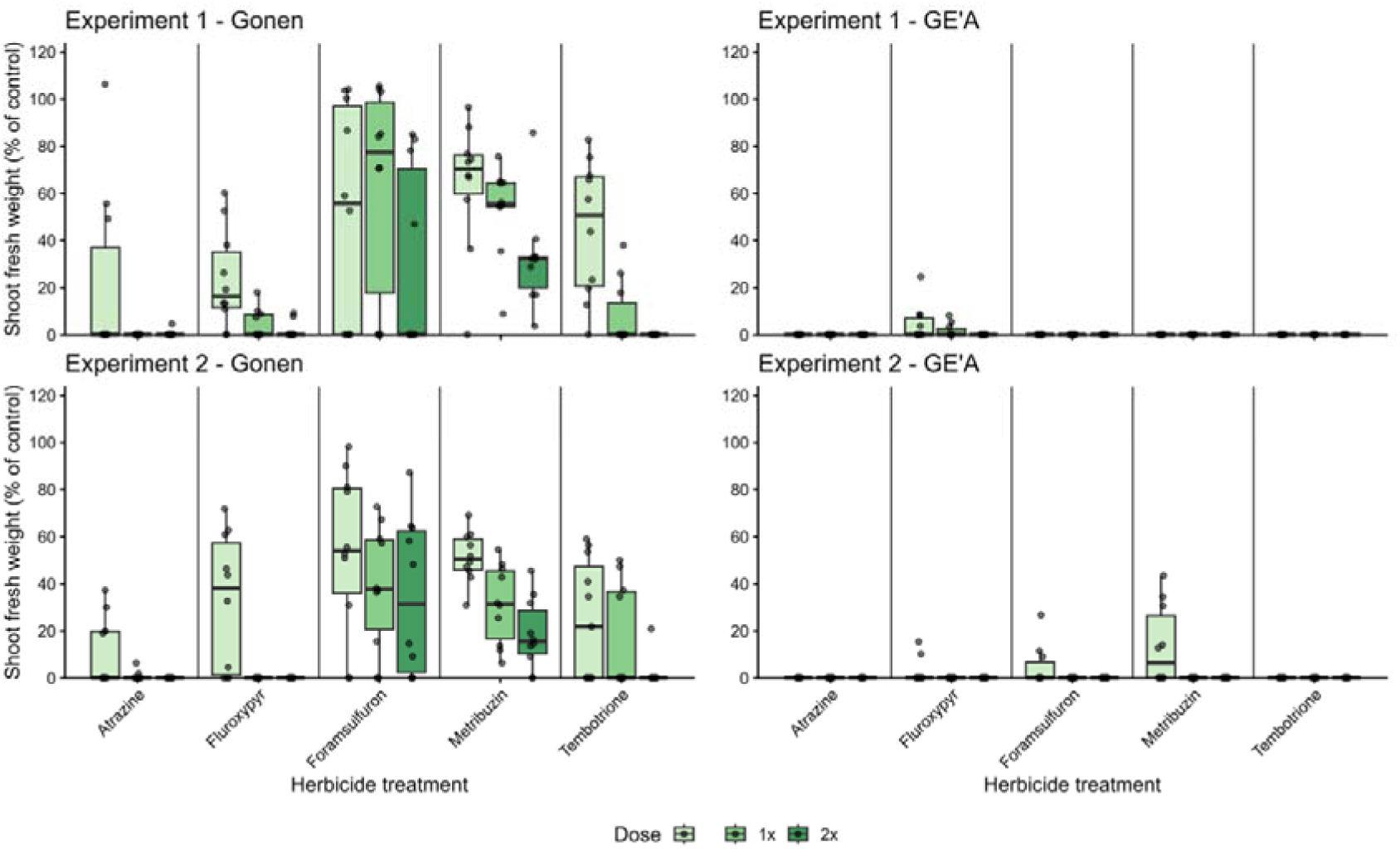
Effect of herbicides from different MOAs on the response of plants from the resistant (Gonen) and sensitive (GE’A) *A. palmeri* populations. Foramsulfuron (ALS - Group 2), fluroxypyr (Auxin Mimic - Group 4), tembotrione (HPPD - Group 27), atrazine and metribuzin (PSII - Group 5). Shoot fresh weight was recorder 21 days after herbicide application.

## 3 RESULTS

### 3.1 Herbicide application

#### 3.1.1 Pre-emergence herbicide application

Pre-emergence herbicides were effective in control of the Gonen population in pot experiments. In the control pots, which were not treated with herbicides, a mean of 5.43±3.69 and 5.86±3.02 plants per pot had emerged and established 29 and 51 DAT, respectively (Table 2). In contrast, pre-emergence application of the HPPD inhibitor isoxaflutole, the VLCFA inhibitor *S*-metolachlor and the combination of fomesefan (PPO inhibitor) and terbutryn (PSII) significantly reduced the emergence of *A. palmeri*, both 29 days and 51 days after the application of the herbicides to the soil, more pronouncedly for *S*-metolachlor (*P*<0.01; Table 2). The reduction in emergence was also reflected in the biomass accumulation. The mean shoot fresh weight was lower than 1% compared to the untreated control at both sampling dates for most herbicide used, excluding the fomesefan + terbutryn treatment were plants had a higher biomass (mean of 0.23 ± 0.39 g; Table 2).

**Table 2.** Plant count and shoot fresh weight (% of control) of the *Amaranthus palmeri* Gonen population treated with pre-emergence herbicides (Dunnett’s post hoc test vs. UTC). Herbicide application was: *S*-metolachlor (VLCFA - Group 15), isoxaflutole (HPPD - Group 27) and a combination fomesafen + terbutryn (PPO - Group 14 + PSII - Group 5).

| Treatment | Sampling date (days after treatment) |  |  |  |
| --- | --- | --- | --- | --- |
|  | 29 days |  | 51 days |  |
| | Mean ± SD | P-value | Mean ± SD | P-value |
| Emergence (No. of plants) |  |  |  |  |
| UTC | 5.43±3.69 | - | 5.86±3.02 | - |
| S-metolachlor | 0.00±0.00 | <0.001 | 0.00±0.00 | <0.001 |
| Isoxaflutole | 1.29±1.89 | 0.008 | 1.57±1.40 | <0.001 |
| Fomesafen + Terbutryn | 1.86±3.02 | 0.024 | 2.00±1.91 | 0.002 |
| Shoot fresh weight (% of control) |  |  |  |  |
| UTC | 2.31±1.22 | - | 0.04±0.04 | - |
| S-metolachlor | 0.00±0.00 | <0.001 | 0.00±0.00 | 0.003 |
| Isoxaflutole | 0.01±0.02 | <0.001 | 0.01 ± 0.02 | 0.031 |
| Fomesafen + Terbutryn | 0.23 ± 0.39 | <0.001 | 0.01 ± 0.01 | 0.031 |

#### 3.1.2 Post-emergence herbicide application

While herbicide application failed to achieve complete control, the magnitude of resistance varied significantly across the tested MOAs, however, the same trend was observed across both experimental runs. In comparison to the sensitive GE’A population, the Gonen population exhibited reduced sensitivity to several commonly used post-emergence herbicides, representing three different MOA’s encompassing ALS, PSII, and HPPD inhibitors (Fig. 1). Plants were shown to be sensitive for the PSII inhibitor, atrazine, as well as the synthetic auxin fluroxypyr. However, for both experimental runs, we have found plants of the Gonen population exhibited high weight when treated with half of the recommended field rate of fluroxypyr in comparison to plants of the sensitive population of GE’A (Exp. 1, 23.46 ± 20.77 vs. 4.63 ± 7.82, Exp. 2, 32.30 ± 28.90 vs. 2.65 ± 5.46, respectively; Table 3). For Gonen population, in Exp.1, 20% survival was recorded under double of the recommended labelled field rate. However, in Exp. 2 none of the plants survived at the recommended labelled field rate.

**Table 3.**
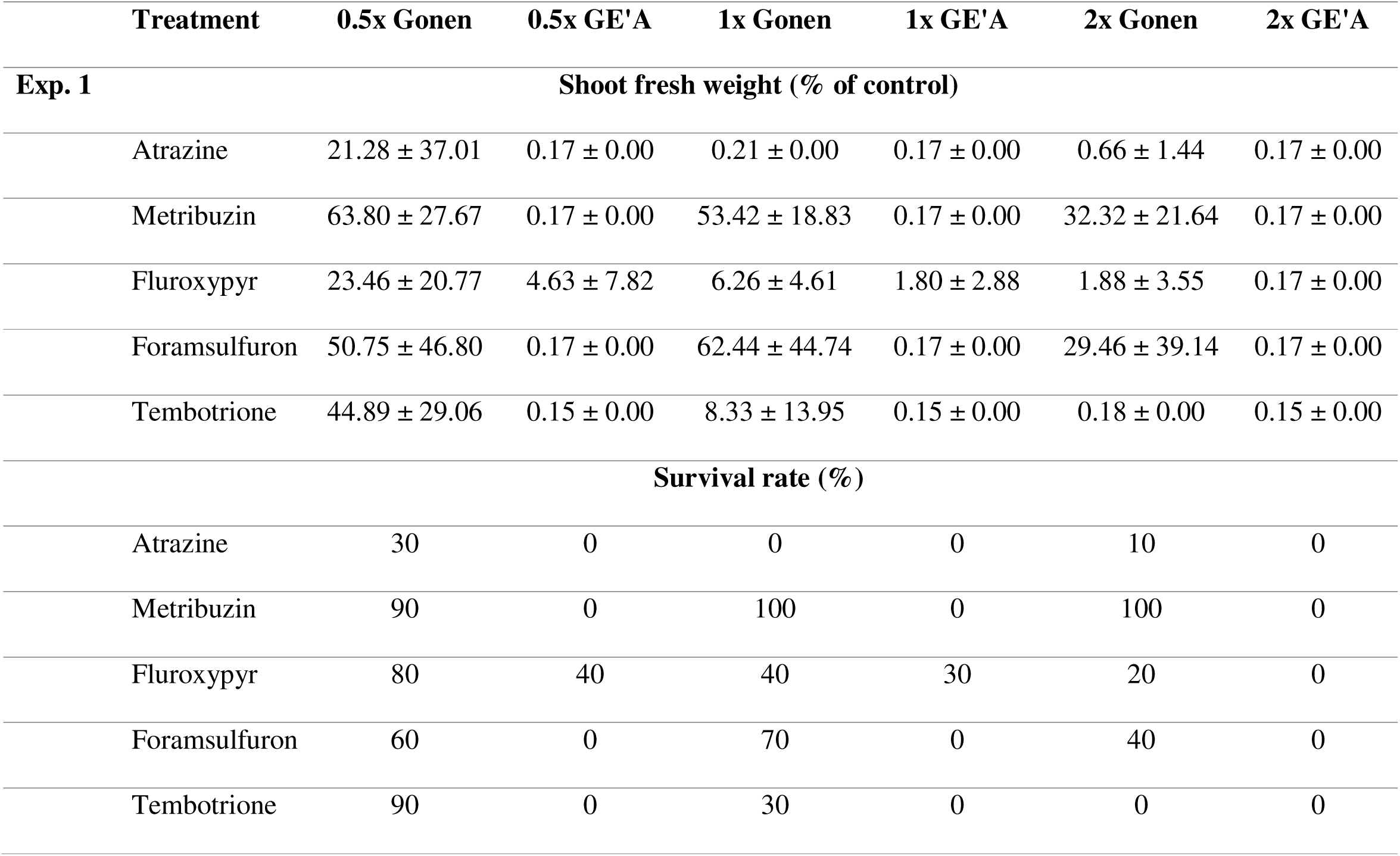
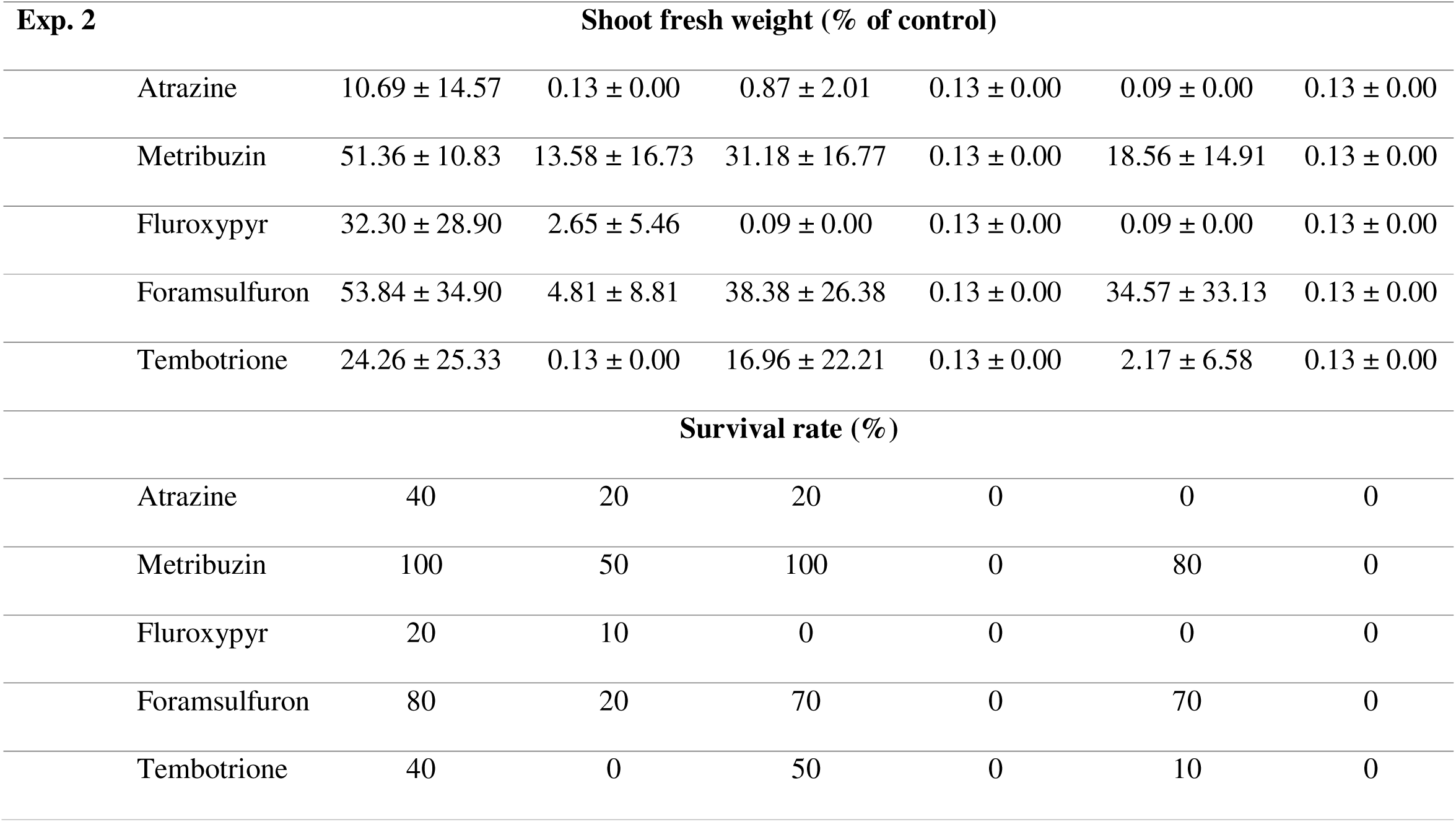
Effect of herbicides from different MOAs on the response of plants from the resistant (Gonen) and sensitive (GE’A) *A. palmeri* populations. Foramsulfuron (ALS - Group 2), Fluroxypyr (Auxin Mimic - Group 4), Tembotrione (HPPD - Group 27), Atrazine and Metribuzin (PSII - Group 5). Shoot fresh weight was recorder 21 days after herbicide application.

|  | Treatment | 0.5x Gonen | 0.5x GE'A | 1x Gonen | 1x GE'A | 2x Gonen | 2x GE'A |
| --- | --- | --- | --- | --- | --- | --- | --- |
| Exp. 1 | Shoot fresh weight (% of control) |  |  |  |  |  |  |
|  | Atrazine | 21.28 ± 37.01 | 0.17 ± 0.00 | 0.21 ± 0.00 | 0.17 ± 0.00 | 0.66 ± 1.44 | 0.17 ± 0.00 |
|  | Metribuzin | 63.80 ± 27.67 | 0.17 ± 0.00 | 53.42 ± 18.83 | 0.17 ± 0.00 | 32.32 ± 21.64 | 0.17 ± 0.00 |
|  | Fluroxypyr | 23.46 ± 20.77 | 4.63 ± 7.82 | 6.26 ± 4.61 | 1.80 ± 2.88 | 1.88 ± 3.55 | 0.17 ± 0.00 |
|  | Foramsulfuron | 50.75 ± 46.80 | 0.17 ± 0.00 | 62.44 ± 44.74 | 0.17 ± 0.00 | 29.46 ± 39.14 | 0.17 ± 0.00 |
|  | Tembotrione | 44.89 ± 29.06 | 0.15 ± 0.00 | 8.33 ± 13.95 | 0.15 ± 0.00 | 0.18 ± 0.00 | 0.15 ± 0.00 |
|  | Survival rate (%) |  |  |  |  |  |  |
|  | Atrazine | 30 | 0 | 0 | 0 | 10 | 0 |
|  | Metribuzin | 90 | 0 | 100 | 0 | 100 | 0 |
|  | Fluroxypyr | 80 | 40 | 40 | 30 | 20 | 0 |
|  | Foramsulfuron | 60 | 0 | 70 | 0 | 40 | 0 |
|  | Tembotrione | 90 | 0 | 30 | 0 | 0 | 0 |

| Exp. 2 | Shoot fresh weight (% of control) |  |  |  |  |  |
| --- | --- | --- | --- | --- | --- | --- |
| Atrazine | 10.69 ± 14.57 | 0.13 ± 0.00 | 0.87 ± 2.01 | 0.13 ± 0.00 | 0.09 ± 0.00 | 0.13 ± 0.00 |
| Metribuzin | 51.36 ± 10.83 | 13.58 ± 16.73 | 31.18 ± 16.77 | 0.13 ± 0.00 | 18.56 ± 14.91 | 0.13 ± 0.00 |
| Fluroxypyr | 32.30 ± 28.90 | 2.65 ± 5.46 | 0.09 ± 0.00 | 0.13 ± 0.00 | 0.09 ± 0.00 | 0.13 ± 0.00 |
| Foramsulfuron | 53.84 ± 34.90 | 4.81 ± 8.81 | 38.38 ± 26.38 | 0.13 ± 0.00 | 34.57 ± 33.13 | 0.13 ± 0.00 |
| Tembotrione | 24.26 ± 25.33 | 0.13 ± 0.00 | 16.96 ± 22.21 | 0.13 ± 0.00 | 2.17 ± 6.58 | 0.13 ± 0.00 |
| Survival rate (%) |  |  |  |  |  |  |
| Atrazine | 40 | 20 | 20 | 0 | 0 | 0 |
| Metribuzin | 100 | 50 | 100 | 0 | 80 | 0 |
| Fluroxypyr | 20 | 10 | 0 | 0 | 0 | 0 |
| Foramsulfuron | 80 | 20 | 70 | 0 | 70 | 0 |
| Tembotrione | 40 | 0 | 50 | 0 | 10 | 0 |

Differential sensitivity was observed between the two PSII inhibitors. In the Gonen population, plants treated with metribuzin (a triazinone) at double the recommended label rate retained substantial shoot fresh weight relative to the untreated control (32.32% ± 21.64 in Exp. 1 and 18.56 ± 14.91 for Exp. 2). Conversely, atrazine provided higher efficacy, at double the recommended rate, plant survival was 10% in Exp. 1 and 0% in Exp. 2, whereas at the single recommended rate, survival reached 20% in Exp. 2 and 0% in Exp. 1 (Table 3). For the sensitive GE’A population, 20% and 50% survival were observed at half the recommended rate for atrazine and metribuzin, respectively, in Exp. 2.

Applications of of oo at double the recommended labelled field rate resulted in reduction in average plant weight in of the Gonen population compared to the untreated control plants (Exp. 1, 29.46 ± 39.14, Exp. 2, 34.57 ± 33.13, respectively; Table 3); however, the survival rate remained at 40% for Exp. 1 and 70% at Exp. 2 (Table 3).

Plants treated with the HPPD inhibitor tembotrione at the recommended labelled filed rate exhibited reduced shoot fresh weight compared to those treated with foramsulfuron and atrazine, with only 8.33 ± 13.95 in Exp. 1 and 16.96 ± 22.21 in Exp. 2. However, survival rate was 30% for Exp. 1 and 50% for Exp. 2, for the later 10% of the plants also survived double of the recommended labelled field rate (Table 3).

### 3.2 Mechanism of resistance

#### 3.2.1 Non-target-site resistance

When evaluating the effect of adding malathion to each herbicide treatment, notable differences in both shoot weight suppression and plant survival emerged between the two populations and across experiments (Table 4).

**Table 4.**
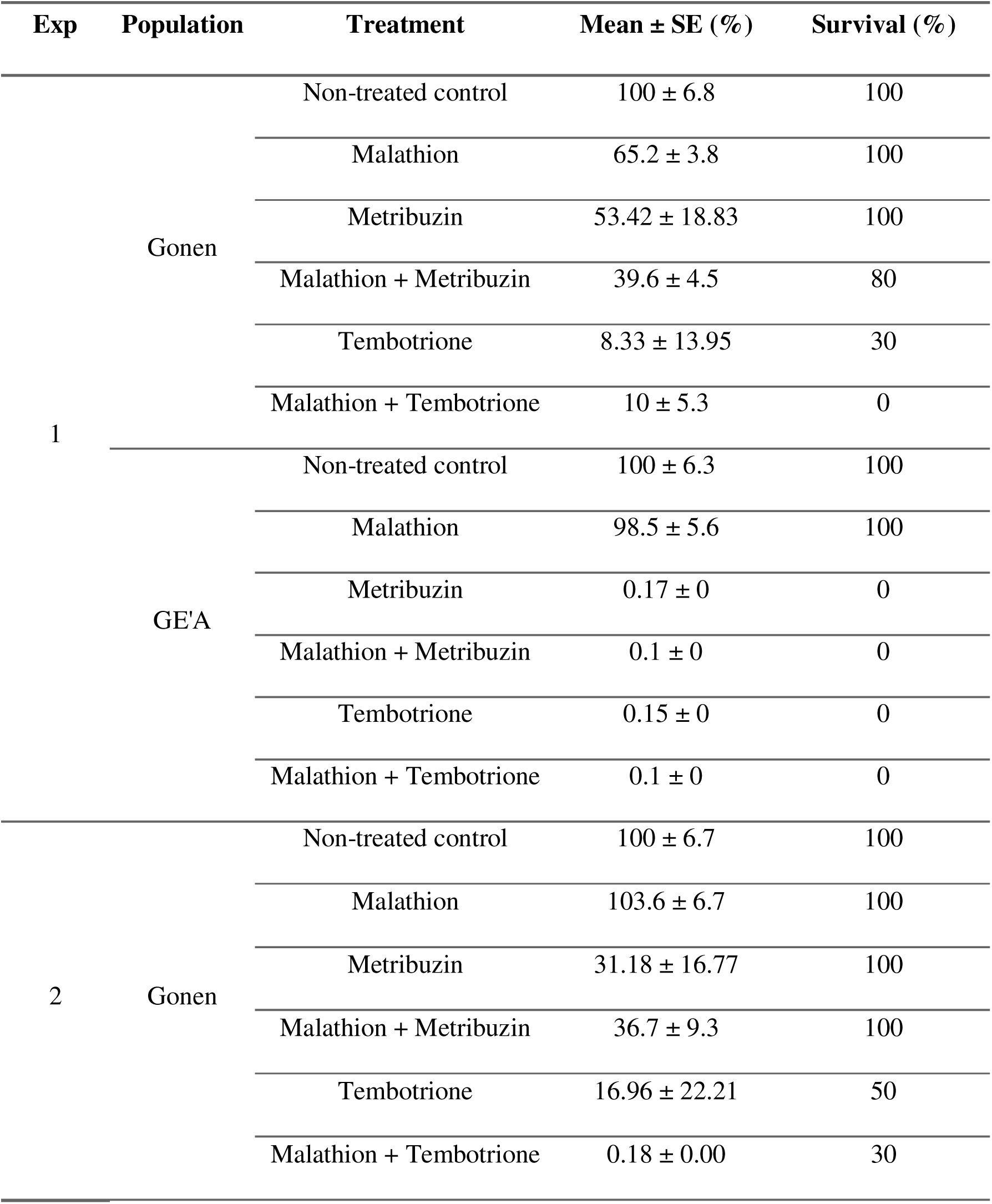
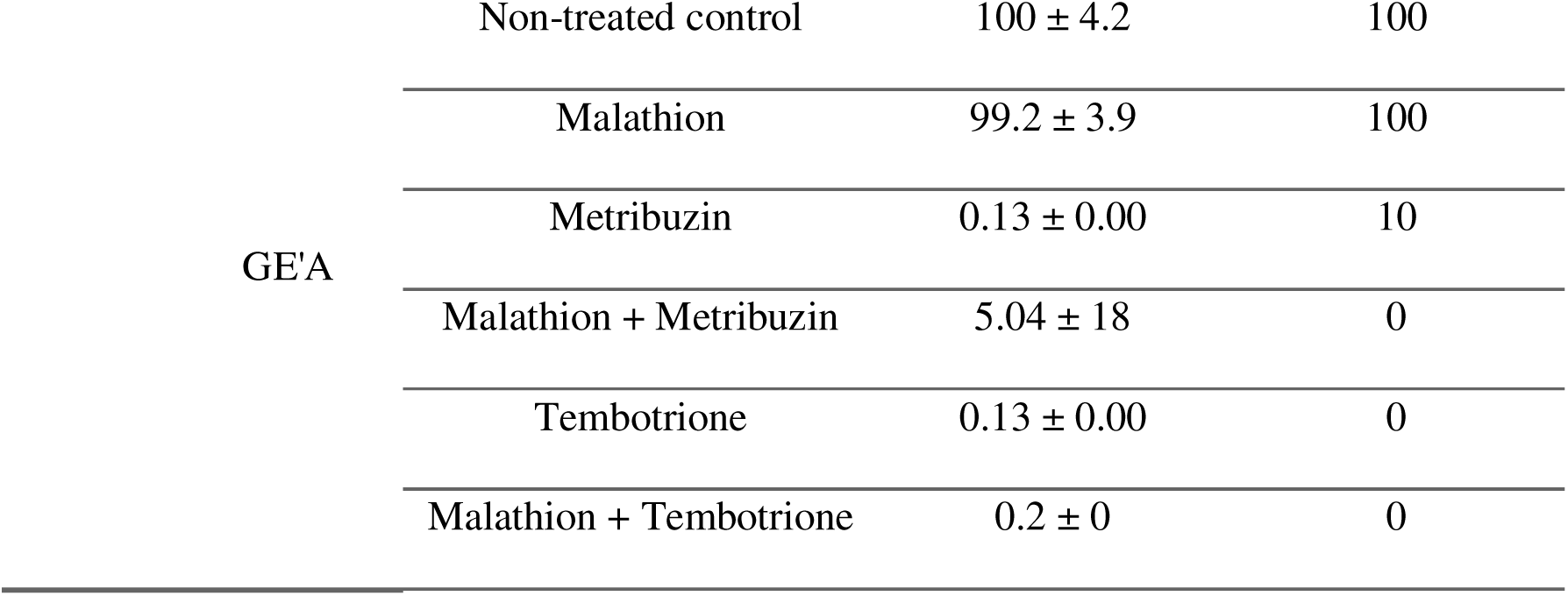
Effect of malathion pre-treatment on the survival rate of plants from resistant (Gonen) and sensitive (GE’A) *A. palmeri* populations. Malathion was applied two hours before the tested herbicide [metribuzin and tembotrione (PSII - Group 5 and HPPD - Group 27, respectively)]. Shoot fresh weight and survival percentage were recorder 21 days after herbicide application.

In the resistant Gonen population, adding malathion to tembotrione consistently enhanced herbicide efficacy, however, effect on average shoot fresh weight was recorded only in Exp. 2 (16.96% down to 0.18%), while survival rate decreased for both experiments, 30% in Exp. 1 (30% to 0%) and 20% in Exp. 2 (50% to 30%) (Table 4). For metribuzin in Gonen, adding malathion caused a marked ∼14% reduction in shoot biomass in Exp. 1 (53.42% down to 39.6%) with slight reduction in survival rate (100% vs. 80%), whereas in Exp. 2 it had minimal impact on biomass (31.18% vs. 36.7%) and no effect on survival rate. Furthermore, for GE’A population, adding malathion to metribuzin successfully eliminated the remaining fraction of surviving plants, reducing survival from 10% to 0% in Exp. 2. Aside from 10% survival in Exp. 2, adding malathion had no effect on the sensitive GE’A population, as all plants were controlled using both herbicide treatments (Table 4).

#### 3.2.2 Target-site resistance to ALS Inhibitors

Target-site resistance to ALS inhibitors is frequently driven by single-nucleotide polymorphisms within the *ALS* gene. Sequence alignment of foramsulfuron-resistant *A. palmeri* individuals from the Gonen population identified a Trp574 to Leu amino acid substitution (Fig. 2A). Four of the seven analysed plants were homozygous (R/R) for this mutant allele. In contrast, target-site sequencing of the *psbA* gene from eight Gonen individuals surviving double the recommended field rate revealed a wild-type sequence across all established resistance hotspots (Fig. 2B), including residues Leu218, Val219, Ala251, Phe255, Ser264, and Phe274.

**Figure 2.**
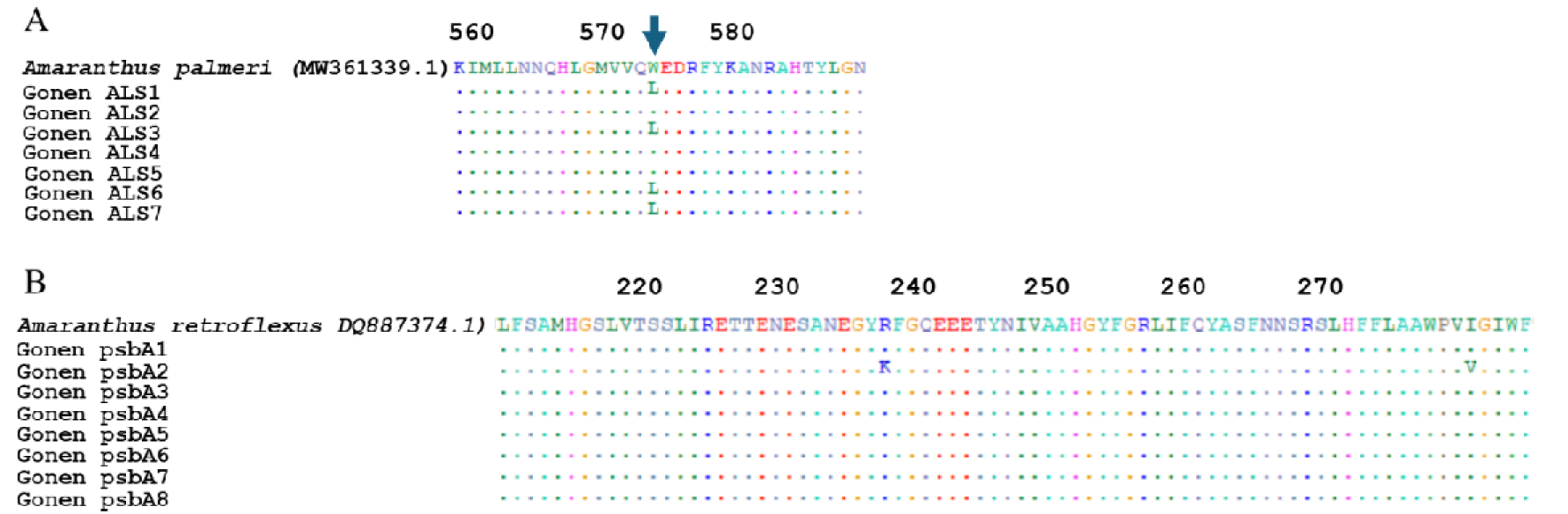
Alignment of partial *ALS* (A) and *psbA* (B) gene sequences from resistant *A. palmeri* plants (n = 7 for *ALS*, n = 8 for *psbA*) originating from Gonen population. Alignments were performed using BioEdit software (Hall, 1999) relative to reference sequences obtained from NCBI GenBank (*A. palmeri ALS*, MW361339.1; *A. retroflexus psbA*, DQ887374.1).

## 4 DISCUSSION

### 4.1 Efficacy of Pre-Emergence Herbicides

In contrast to post-emergence failures, the Gonen population exhibited complete susceptibility to pre-emergence herbicide applications, maintaining robust control for at least 51 days post-application. With the exception of isoxaflutole, all evaluated pre-emergence chemistries are moderately persistent in soil (Tzilivakis et al., 2026). Although parent isoxaflutole degrades rapidly, its primary active metabolite, diketonitrile, possesses higher soil persistence that extends residual activity (Papiernik et al., 2007). Sustained residual control confirms that field-level failures are driven by multi-MOA post-emergence resistance rather than rapid herbicide degradation in soil.

Specifically, susceptibility to VLCFA-inhibiting (*S*-metolachlor) and PPO-inhibiting (fomesafen) herbicides provides immediate management options for this multi-resistant biotype. The complete control observed with fomesafen + terbutryn tank mixes was likely driven primarily by fomesafen, however, seedling vulnerability may have enabled terbutryn activity despite post-emergence PSII resistance. Non-target-site metabolic resistance is often phenology-dependent, leaving emerging seedlings significantly more vulnerable than established plants (Riechers et al., 2024). This developmental vulnerability is further illustrated by isoxaflutole, which provided complete control pre-emergence despite poor post-emergence control with tembotrione (HPPD, same MOA).

### 4.2 Post-emergence herbicide application and resistance mechanisms

Evaluating the Gonen population across four major herbicide MOAs revealed significant post-emergence control failures compared to the susceptible GE’A population. For the ALS inhibitor foramsulfuron, Gonen plants demonstrated 40% survival in the first experimental run and 70% in the second under double the recommended labelled field rate. Resistance to ALS inhibitors is well-established regionally, trifloxysulfuron resistance has been documented in several Israeli biotypes, including populations from the same agricultural region (Abu-Nassar et al., 2024). Similar ALS-resistance profiles driven by target-site mutations (e.g., Trp574 to Leu, Pro197 to Thr, Asp376 to Glu) occur across Europe and the Mediterranean basin (Manicardi et al., 2023), as well as in Turkey, where populations exhibit cross-resistance to foramsulfuron and nicosulfuron alongside glyphosate (EPSPS) resistance (Kaya-Altop et al., 2025). Previous studies showed that Trp574 to Leu confer high level of resistance to broad spectrum of ALS inhibitors in *A. palmeri* (Larran et al., 2017; Singh et al., 2019). It is reasonable to hypothesize that this mutation also induces cross-resistance to other untested ALS-inhibiting herbicides.

Differential sensitivity was observed within PSII inhibitors. Metribuzin (a triazinone) resulted in virtually no reduction in plant survival under twice the recommended labelled field rates, whereas atrazine (a triazine) provided high control efficacy. Sequencing of eight individuals surviving double the recommended rate of metribuzin revealed no mutations in the *psbA* gene (including the Ala251 to Val mutation, which typically confers triazinone resistance while preserving triazine sensitivity; Gaines et al., 2020). Furthermore, pre-application of the cytochrome P450 inhibitor malathion failed to reverse metribuzin resistance, ruling out both standard P450-mediated metabolism and direct *psbA* target-site mutations, pointing to alternative non-target-site mechanisms.

Tembotrione treatments yielded 30-50% survival at the recommended field rate and up to 10% survival at double the field rate. The surviving phenotype under tembotrione strongly suggests non-target-site metabolic resistance, with the use of malathion. In *A. palmeri*, HPPD resistance is frequently driven by enhanced P450-catalyzed detoxification, specifically linked to the upregulated expression of genes such as *CYP81E8* (Shyam et al., 2022) and *CYP72A1182* (Rigon et al., 2025). Differences between experimental runs can be attributes to variation in environmental conditions. Differences between experimental runs can be attributed to variations in environmental conditions. Previous studies have demonstrated that malathion efficacy is temperature-sensitive, specifically, elevated temperatures post-application can reduce its capacity to inhibit herbicide detoxification (Matzrafi et al 2016).

As resistance to traditional auxins like dicamba and 2,4-D increases globally, growers have increasingly shifted to pyridine-carboxylic acids like fluroxypyr, which may retain efficacy due to distinct molecular binding or metabolic profiles (Busi et al., 2018; Kumar et al., 2019). For the synthetic auxin fluroxypyr, 40% of the Gonen population and 30% of the GE’A population survived the labelled field rate in the initial run. However, surviving plants exhibited severe physiological distortion, growing twisted and close to the pot surface, rendering them non-competitive against a maize crop. In the subsequent run (Exp. 2), no plants survived the labelled rate. The transient survival and phenotype observed across both biotypes suggest strong environmental modulation of fluroxypyr efficacy rather than stable genetic resistance.

The high-level resistance observed in the Gonen population to the primary herbicides utilized in Israeli maize production presents a critical challenge for regional weed management. Overall findings confirm multi-MOA resistance across major post-emergence herbicides, while highlighting soil-applied, residual pre-emergence applications as an essential foundation for effective control. Stacked multiple resistance across five to six herbicide modes of action has been documented in North American *Amaranthus* species, such as *A. tuberculatus* in Missouri (Shergill et al., 2018) and *A. palmeri* in Kansas (Shyam et al., 2021). Similar to the Gonen population, biotypes like the Kansas KCTR population have evolved stacked resistance by combining target-site mutations (e.g., Trp574 to Leu in the *ALS* gene) with non-target-site metabolic mechanisms that confer cross-resistance to HPPD and PSII inhibitors.

## CONCLUSIONS

Future weed management strategies in Israeli maize must minimize sole reliance on post-emergence foliar applications and prioritize residual pre-emergence programs to reduce selection pressure. Integrating non-chemical strategies, such as strategic tillage, crop rotation, and competitive cover crops, will be essential to exhaust the *A. palmeri* seed bank and maintain crop production sustainability in the region.

## ACKNOWLEDGEMENTS

The authors thank Eran Kenan and On Rabinovitch for their assistance with field collection of seed and soil samples.

